# Development and Validation of Phenotype Classifiers across Multiple Sites in the Observational Health Sciences and Informatics (OHDSI) Network

**DOI:** 10.1101/673418

**Authors:** Mehr Kashyap, Martin Seneviratne, Juan M Banda, Thomas Falconer, Borim Ryu, Sooyoung Yoo, George Hripcsak, Nigam H Shah

## Abstract

**Objective:** Accurate electronic phenotyping is essential to support collaborative observational research. Supervised machine learning methods can be used to train phenotype classifiers in a high-throughput manner using imperfectly labeled data. We developed ten phenotype classifiers using this approach and evaluated performance across multiple sites within the Observational Health Sciences and Informatics (OHDSI) network.

**Materials and Methods:** We constructed classifiers using the Automated PHenotype Routine for Observational Definition, Identification, Training and Evaluation (APHRODITE) R-package, an open-source framework for learning phenotype classifiers using datasets in the OMOP CDM. We labeled training data based on the presence of multiple mentions of disease-specific codes. Performance was evaluated on cohorts derived using rule-based definitions and real-world disease prevalence. Classifiers were developed and evaluated across three medical centers, including one international site.

**Results:** Compared to the multiple mentions labeling heuristic, classifiers showed a mean recall boost of 0.43 with a mean precision loss of 0.17. Performance decreased slightly when classifiers were shared across medical centers, with mean recall and precision decreasing by 0.08 and 0.01, respectively, at a site within the USA, and by 0.18 and 0.10, respectively, at an international site.

**Discussion and Conclusion:** We demonstrate a high-throughput pipeline for constructing and sharing phenotype classifiers across multiple sites within the OHDSI network using APHRODITE. Classifiers exhibit good portability between sites within the USA, however limited portability internationally, indicating that classifier generalizability may have geographic limitations, and consequently, sharing the classifier-building recipe, rather than the pre-trained classifiers, may be more useful for facilitating collaborative observational research.

## BACKGROUND AND SIGNIFICANCE

Electronic phenotyping refers to the task of identifying patients within an electronic health record (EHR) who match a defined clinical profile [1]. Accurate phenotyping is critical to support observational research, pragmatic clinical trials, quality improvement evaluations, and clinical decision support systems [2,3]. However, issues such as missingness, accuracy and heterogeneity in EHR data present major challenges to effective phenotyping [4].

The traditional approach to phenotyping has been rule-based, where a cohort is manually defined with inclusion and exclusion criteria based on structured data such as diagnosis codes, laboratory results and medications [5]. Although several collaborative networks exist for generating and sharing rule-based definitions, including the PheKB (Phenotype Knowledge Base) [6], these phenotypes are typically labor-intensive to create and require multiple rounds of review by domain experts [7].

Recent efforts to establish common data models for EHRs, including the Observational Health Data Sciences and Informatics (OHDSI) [8] and the Informatics for Integrating Biology and the Bedside (i2b2) initiatives [9], are enabling large-scale observational research and algorithm deployment across sites. To make use of this infrastructure, we need the ability to generate complex, generalizable phenotypes more rapidly than rule-based approaches allow [3].

Supervised machine learning has emerged as a way to generate phenotypes in a high-throughput manner [1]. By incorporating a wide range of EHR features, statistical methods have shown robust performance for complex phenotypes including chronic pain and rheumatoid arthritis [10,11] with some evidence to indicate portability (preserved classification accuracy) across sites [12]. The major bottleneck for supervised machine learning is access to labeled training data, which traditionally requires manual chart review by clinicians.

To address the scarcity of labeled training data, Chen *et al.* used active learning to intelligently select training samples for labeling, demonstrating that classifier performance could be preserved with fewer samples [13]. Another trend is the use of “silver standard training sets”, a semi-supervised approach where training samples are labeled using an imperfect heuristic rather than by manual review [14–19]. The intuition is that noise-tolerant classifiers trained on imperfectly labeled data will abstract higher order properties of the phenotype beyond the original labeling heuristic (so-called noise-tolerant learning [20]). Halpern *et al.* have described the anchor learning framework where the presence of ‘anchor’ references, which are highly predictive of a phenotype and are conditionally independent of other features (i.e. best predicted by the phenotype itself), are used to define an imperfect training cohort for phenotype classifiers [16]. Similarly, Agarwal *et al.* developed the XPRESS (eXtraction of Phenotypes from Records using Silver Standards) pipeline where noisy training samples are defined based on highly specific keyword mentions in a patient’s EHR [14]. This led to the development of the Automated PHenotype Routine for Observational Definition, Identification, Training and Evaluation (APHRODITE) R-package, an open-source implementation of the XPRESS framework with dynamic anchor learning built on the OHDSI common data model, which has shown comparable performance to rule-based definitions for two phenotypes (type 2 diabetes and myocardial infarction) [21].

The current work addresses two questions resulting from the use of APHRODITE. The first is about the labeling function used to generate imperfectly labeled training data. While APHRODITE uses the mention of a single phrase, we hypothesize that a high-precision labeling heuristic based on multiple keyword (or phrase) mentions may improve classifier performance in situations where phenotyping precision is critical. In addition, APHRODITE was evaluated using balanced cohorts of cases and controls; in real-world situations where the number of controls far outnumbers cases, a higher-precision labeling function may perform better. We investigate the improvement obtained via complex labeling functions across a spectrum of ten different phenotypes, and using real-world disease prevalence in the test data.

The second question is about APHRODITE’s ability to port both the final classifiers and the underlying training ‘recipes’ between OHDSI sites. A recent study demonstrated the translation of PheKB definitions into executable EHR queries that ported across six different health systems [22]; however the portability of classifier-based approaches such as APHRODITE has yet to be rigorously assessed. We conduct reciprocal experiments where we evaluate the performance of phenotype classifiers trained at our academic medical center on the EHRs of two other health systems, and conversely, evaluate the performance of classifiers trained externally on our data. We find that phenotype classifiers perform well across OHDSI sites, though portability may be limited by underlying differences in EHR data at each site.

## MATERIALS AND METHODS

### Data Sources

We used longitudinal electronic health record data from Stanford Hospital & Clinics and Lucile Packard Children’s Hospital, Columbia University Medical Center, and Seoul National University Bundang Hospital (SNUBH) to construct and evaluate phenotype classifiers. At Stanford, patient data was extracted from the Stanford Medicine Research Data Repository clinical data warehouse and included nearly 1.8 million patients and 53 million unique visits. The dataset used at Columbia comprised of over 5 million patients from the New York Presbyterian Hospital clinical data warehouse. At SNUBH, the EHR dataset included over 1.8 million patients. Patient data at each institution was composed of coded diagnoses, laboratory tests, medication orders, and procedures. All data at the three institutions were mapped to the Observational Medical Outcomes Partnership (OMOP) Common Data Model (CDM), which serves as a shared standard representation of clinical data across multiple data sources and institutions.

### Phenotype selection and classifier development

We selected ten phenotypes (appendicitis, type 2 diabetes mellitus, cataracts, heart failure, abdominal aortic aneurysm, epileptic seizure, peripheral arterial disease, adult onset obesity, glaucoma, and venous thromboembolism) for which rule-based definitions have been created by either the eMERGE or OHDSI networks. We developed classifiers for each phenotype using the APHRODITE framework, an R-package built for the OMOP CDM that can be used to construct phenotype classifiers using imperfectly labeled training data. In previous work, the labeling heuristic used with APHRODITE was based on single mentions of relevant terms in textual data. In this study, we used multiple mentions of disease-specific codes as our labeling function. In particular, we identified cases by searching patients’ clinical data for at least four mentions of any relevant SNOMED code associated with the phenotype of interest (Figure 1). We identified all relevant codes by using vocabulary tables and existing relationships between concepts within the OMOP CDM. Patients who did not meet this multiple mention criteria were considered controls for training purposes, and the ratio of training cases to controls was set to 1:1. We used four mentions since we expected this to be a highly specific label for any phenotype of interest. This number was also low enough such that it most consistently yielded an adequate number of cases (250+ based on experiments not reported in this study) to train classifiers. When sufficient cases were not identified, we incrementally lowered the number of mentions to identify more cases.

Once the training cohort was identified, we represented patient data with the following feature types: visits, observations, lab results, procedures and drug exposures. Frequency counts were calculated for each feature capturing the entire course of patients’ EHR records. We chose not to exclude a single mention of relevant disease-specific codes as potential features used by classifiers, since our labeling function was based on multiple mentions. Random forest classifiers were trained for each phenotype using 5-fold cross validation.

**Figure 1.**
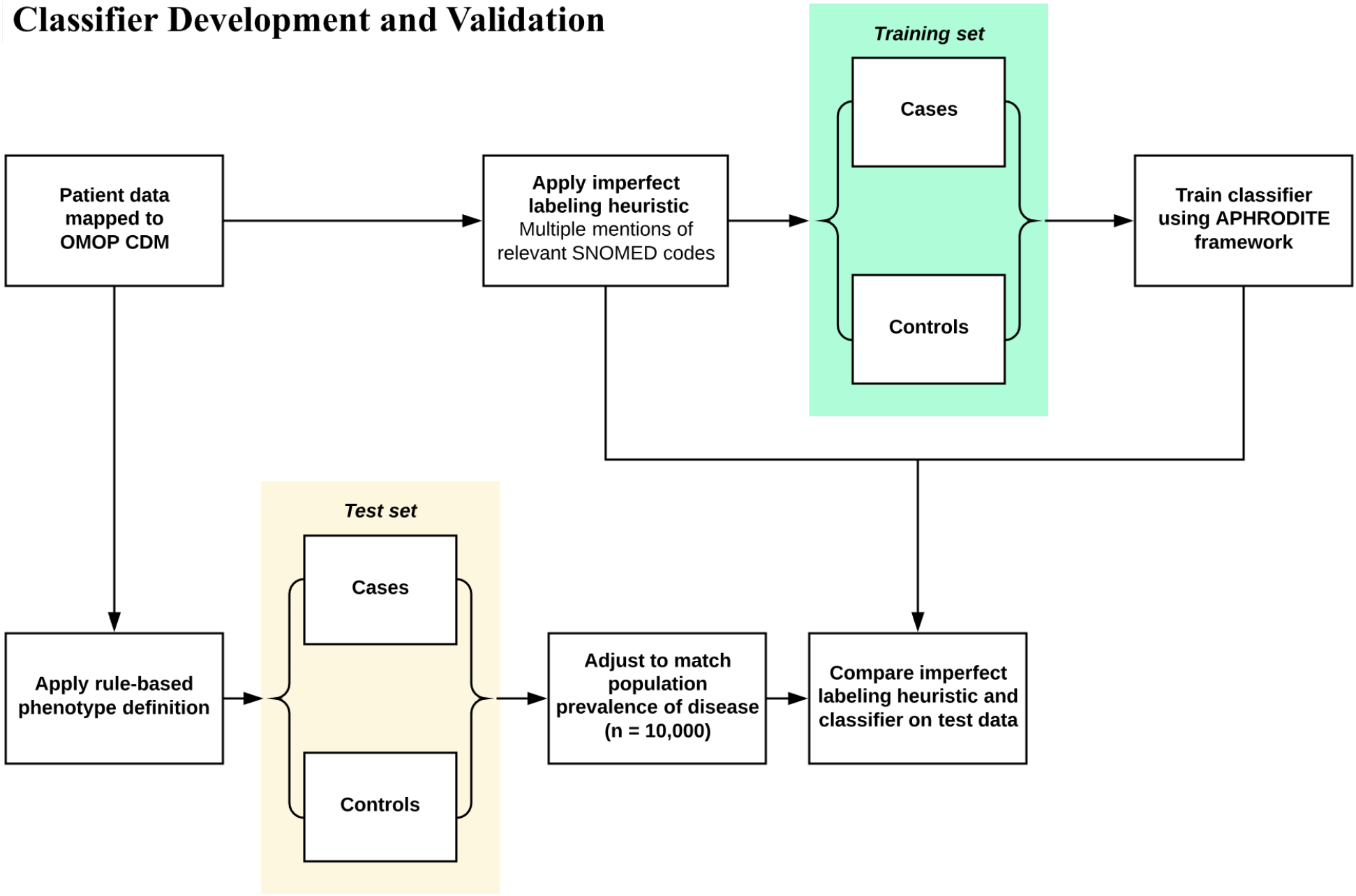
Development and validation of phenotype classifiers. Training sets were constructed by applying multiple-mentions based imperfect labeling function to our patient data extract. Patients with multiple mentions of any SNOMED codes relevant to the phenotype of interest were considered training cases. Patients who did not meet this criteria were labeled as training controls. Random forest classifiers were built for each phenotype using 5-fold cross validation. The test set was constructed using OMOP implementations of rule-based phenotype definitions. Test cases were randomly sampled from the cohort of patients selected by the rule-based definitions. Test controls were sampled from the remaining patients. For each phenotype, the imperfect labeling function used to generate the training set and the corresponding classifier were evaluated using the rule-based phenotype derived test sets.

### Classifier validation with cohorts derived from rule-based definitions

#### Development of evaluation sets

Rule-based definitions were used to identify the cohort of patients comprising the test set for each phenotype (Figure 1). Two of the definitions, appendicitis and cataracts, were OMOP implementations of definitions that were publicly available on Phenotype KnowledgeBase (PheKB), a repository of phenotype algorithms developed by the eMERGE network. The other eight definitions were developed and evaluated collaboratively by several members of the OHDSI network with clinician oversight. Although PheKB definitions have been shown to favor precision and have low recall relative to manual chart review, these rule-based definitions were the best available ground truth label for this experiment.

Rule-based definitions were implemented using ATLAS, an open source software tool for building patient cohorts with OMOP CDM mapped data. Test cases were identified by randomly sampling the cohort of patients selected by the rule-based definitions. Test controls were identified by randomly sampling from the remaining patients. All test sets were composed of 10,000 patients, with the proportion of cases set equal to the population prevalence of the corresponding phenotype. Any patients used to train classifiers were excluded from test sets.

#### Local validation of phenotype classifiers

We evaluated the performance of our classifiers by running them on the test sets derived from our rule-based definitions. Classifiers were evaluated locally by using our patient data extract. For reference, we also assessed the performance of the ‘multiple mentions’ labeling heuristic described previously. Performance was reported in terms of recall and precision.

### Performance of classifiers across multiple sites

To evaluate the portability of our phenotype classifiers, we shared the ten classifiers developed on our patient data extract with two other institutions within the OHDSI network, Columbia University and SNUBH. Since both these institutions have mapped their patient datasets to meet OMOP CDM specifications, we were able to share our classifiers without any modification. Classifier performance was evaluated in a process identical to the one used locally – at both sites, rule-based definitions were used to derive the cohort of patients comprising the test set for each phenotype.

We further assessed the portability of phenotype classifiers by performing a reciprocal experiment in which models were constructed at Columbia and SNUBH, and subsequently evaluated on our patient data extract. Classifiers were developed for all ten phenotypes using the same method that was used locally. Specifically, we employed the same labeling approach to generate training sets for each phenotype – patients with multiple mentions of relevant disease-specific codes were considered training cases, while others were considered training controls. Once classifiers were developed at Columbia and SNUBH, we evaluated their performance on our patient data extract, using test sets that were constructed from rule-based definitions.

### Comparing demographics of cases across sites

While rule-based definitions offer an alternative to manual chart review for the generation of test sets, development of these definitions is ultimately labor-intensive and limits the speed with which classifiers can be evaluated. To circumvent the need for rule-based definitions and chart review, we propose comparing the demographics of patients identified as cases by our classifiers across different sites as a proxy for model validation. For this experiment, we randomly selected 150,000 patients at each site and used classifiers developed at our institution to identify cases for each phenotype. We then compared the cohorts of patients labeled as cases across different sites with respect to key demographics, including age and sex. The purpose of this was to evaluate whether the classifiers not only showed comparable performance across sites, but also identified comparable cohorts in terms of basic demographic features.

## RESULTS

### Local performance of classifiers

We first compared the performance of our phenotype classifiers with the ‘multiple mentions’ labeling heuristic used to identify training cases for each phenotype. Table 1 shows the recall and precision of both of these phenotyping approaches. Requiring multiple disease-specific code mentions to classify patients as cases yields a mean precision of 0.99, as it is likely that patients with several mentions of a relevant code have the associated phenotype. Achieving high precision, however, results in noticeably low recall. The mean recall for requiring multiple mentions was 0.17.

**Table 1.** Test set performance of labeling heuristic requiring multiple disease-specific code mentions compared to phenotype classifiers trained with data labeled using this multiple mentions approach. T2DM = type 2 diabetes mellitus, HF = heart failure, AAA = abdominal aortic aneurysm, PAD = peripheral arterial disease, VTE = venous thromboembolism

| Phenotype | Prevalence of cases in test set | Multiple mentions of SNOMED code |  |  | APHRODITE classifier |  | Recall boost using classifier | Precision loss using classifier |
| --- | --- | --- | --- | --- | --- | --- | --- | --- |
|  |  | No. of mentions | Recall | Precision | Recall | Precision |  |  |
| Appendicitis | 0.05 | 2 | 0.31 | 1.00 | 0.97 | 0.99 | **+0.66** | **-0.01** |
| T2DM | 0.14 | 4 | 0.24 | 0.99 | 0.60 | 0.91 | **+0.36** | **-0.08** |
| Cataracts | 0.17 | 4 | 0.07 | 0.97 | 0.63 | 0.93 | **+0.56** | **-0.04** |
| HF | 0.02 | 4 | 0.33 | 0.94 | 0.99 | 0.56 | **+0.66** | **-0.38** |
| AAA | 0.04 | 4 | 0.22 | 0.99 | 0.53 | 0.97 | **+0.31** | **-0.02** |
| Epileptic seizure | 0.02 | 4 | 0.06 | 1.00 | 0.22 | 0.94 | **+0.17** | **-0.06** |
| PAD | 0.05 | 4 | 0.18 | 0.98 | 0.91 | 0.91 | **+0.72** | **-0.07** |
| Adult onset obesity | 0.36 | 4 | 0.20 | 1.00 | 0.29 | 0.91 | **+0.09** | **-0.09** |
| Glaucoma | 0.01 | 4 | 0.08 | 1.00 | 0.22 | 0.88 | **+0.14** | **-0.12** |
| VTE | 0.01 | 4 | 0.03 | 1.00 | 0.69 | 0.22 | **+0.66** | **-0.78** |

Classifiers built with training data labeled using the multiple code mentions heuristic showed markedly improved recall with relatively small losses in precision. The mean recall boost observed was 0.43 while the mean precision loss was 0.17. Seven classifiers showed precision losses that were less than 0.10. Classifiers for two phenotypes, heart failure and venous thromboembolism, had more considerable losses in precision (−0.38 and −0.78, respectively).

### Performance of classifiers across sites

We evaluated the portability of our classifiers by assessing their performance at Columbia and SNUBH. Table 2 summarizes performance at these two sites. When classifiers were tested at Columbia, mean recall and precision decreased marginally by 0.08 and 0.01, respectively, compared to local performance. Classifiers tested at SNUBH had more significant losses in performance. Mean recall and precision decreased by 0.18 and 0.10, respectively.

**Table 2.** Classifier performance at three sites within OHDSI network. Phenotype classifiers constructed at Stanford were shared with Columbia and SNUBH, and evaluated using test sets derived locally at each site using rule-based definitions. Furthermore, classifiers built at Columbia and SNUBH were shared with Stanford and evaluated using similarly constructed test sets. Blue denotes values equal to 1, white denotes values equal to 0.

| Development site | Stanford |  |  | Columbia | SNUBH |  | Stanford |  |  | Columbia | SNUBH |  |
| --- | --- | --- | --- | --- | --- | --- | --- | --- | --- | --- | --- | --- |
| Validation site | Stanford | Columbia | SNUBH | Stanford | Stanford | SNUBH | Stanford | Columbia | SNUBH | Stanford | Stanford | SNUBH |
| Phenotype | Recall |  |  |  |  |  | Precision |  |  |  |  |  |
| Appendicitis | 0.97 | 0.9 | 0.09 | 0.82 | 0.52 | 0.1 | 0.99 | 0.9 | 0.56 | 0.83 | 0.13 | 0.98 |
| T2DM | 0.6 | 0.63 | 0.77 | 0.58 | 0.67 | 0.75 | 0.91 | 0.86 | 0.75 | 0.75 | 0.51 | 0.89 |
| Cataracts | 0.63 | 0.45 | 0.84 | 0.8 | 0.35 | 0.84 | 0.93 | 0.79 | 0.85 | 0.74 | 0.42 | 0.74 |
| HF | 0.99 | 0.97 | 0.8 | 0.99 | 0.71 | 0.82 | 0.56 | 0.67 | 0.75 | 0.47 | 0.11 | 0.66 |
| AAA | 0.53 | 0.24 | 0.54 | 0.59 | 0.33 | 0.57 | 0.97 | 0.75 | 0.87 | 0.96 | 0.13 | 0.47 |
| Epileptic seizure | 0.22 | 0.3 | 0.28 | 0.41 | 0.46 | 0.11 | 0.94 | 0.87 | 0.55 | 0.79 | 0.08 | 0.68 |
| PAD | 0.91 | 0.89 | 0.57 | 0.48 | 0.46 | 0.55 | 0.91 | 0.87 | 0.68 | 0.69 | 0.24 | 0.59 |
| Adult onset obesity | 0.29 | 0.33 | 0.07 | 0.14 | 0.39 | 0.07 | 0.91 | 0.93 | 0.73 | 0.85 | 0.68 | 0.8 |
| Glaucoma | 0.22 | 0.18 | 0.11 | 0.34 | 0.22 | 0.12 | 0.88 | 0.78 | 0.65 | 0.69 | 0.06 | 0.75 |
| VTE | 0.69 | 0.34 | 0.2 | 0.21 | 0.46 | 0.19 | 0.2 | 0.71 | 0.83 | 0.51 | 0.05 | 0.78 |

We further assessed the portability of classifiers by constructing models at Columbia and SNUBH, and evaluating their performance on our patient data extract. Classifiers developed at Columbia had comparable performance to those developed at Stanford. Specifically, classifiers demonstrated high precision when tested at Stanford, with these classifiers having a mean precision value of 0.73. Mean recall for these classifiers was 0.54. In contrast, classifiers developed at SNUBH did not port as well. For these classifiers, we observed generally worse performance. Mean recall and precision for these classifiers was 0.46 and 0.24, respectively.

### Classifier evaluation by comparing demographic features of cases across sites

We further examined classifier performance by evaluating the demographics of patients classified as cases for each phenotype at all three sites. We used the classifiers built at Stanford to select cases at all sites. The aim of this is to assess both the overall performance and portability of classifiers by determining whether classifiers identify comparable cohorts of patients across sites. Overall, age and proportion of each sex were similar among all patients at the three sites – mean age was 39.3, 39.9, and 40.1 and proportion of males was 0.45, 0.45. and 0.48 at Stanford, Columbia and SNUBH, respectively. However, there was considerable variability in demographics of patients selected as cases by classifiers for each phenotype. For instance, for seven of the ten phenotypes, there was a statistically significant difference in the proportion of males identified as cases. Similar variation existed with regards to mean age of cases at the three sites (supplemental table 1).

## DISCUSSION

This study outlines a method for generating high-precision phenotyping classifiers in a semi-supervised manner. We demonstrate that classifiers trained using a high-precision labeling heuristic (i.e. multiple mentions of disease-specific codes) are able to preserve precision while boosting recall relative to the original labeling function. This recall boost was observed across all ten phenotypes. Furthermore, these classifiers are significantly faster to generate than rule-based phenotype definitions, and do not rely on expert clinical input. This may be a template for high-throughput [3] creation of phenotyping classifiers in a way that optimizes precision and recall, which would greatly facilitate observational research.

An important advantage of building phenotype classifiers with the APHRODITE framework is the ability to easily exchange models across sites. We evaluated model portability in this study by sharing our phenotype classifiers with Columbia and SNUBH. Performance at both of these sites was generally good, with minimal losses in recall and precision at Columbia (0.08 and 0.01, respectively) and a larger performance drop at SNUBH (0.18 and 0.10, respectively). We suspect that the larger drop in performance at SNUBH, which is an international site, is likely related to geographic differences in EHR data and how diagnoses are coded.

In a reciprocal experiment, both Columbia and SNUBH constructed classifiers at their sites and we evaluated these at Stanford. Classifiers developed at Columbia performed well, with these classifiers having a mean precision value of 0.73. Unlike classifiers constructed at Columbia, those developed at SNUBH did not port well, with mean recall and precision for these classifiers of 0.46 and 0.24, respectively.

The poor portability of classifiers developed at SNUBH suggests that in certain cases, in which there are significant differences in the characteristics of the underlying EHR data, sharing the classifier-building recipe (i.e. high precision labeling function), rather than the pre-trained classifiers, may prove more useful. The APHRODITE framework specifically offers the ability to exchange recipes. Unlike traditional supervised learning approaches for phenotyping which require manually searching for patients to construct the training set, sharing a high precision labeling function for generating a large imperfectly labeled training set and re-building classifiers at any given site is efficient and feasible.

This study was limited by the use of PheKB definitions as the gold standard for classifier evaluation. Although these definitions have been reviewed by clinical experts, they are still rule-based definitions with imperfect classification accuracy. While the use of these definitions provided a standardized way to assign ‘ground-truth’ labels across multiple international sites, in future our classifier pipeline could be assessed against clinician-labeled test-sets at each site. The feature engineering scheme used in training classifiers is relatively rudimentary – simply a frequency count of all structured data elements. The performance of our classifiers may therefore be seen as a conservative estimate of such semi-supervised learning. More sophisticated feature engineering regimens such as EHR embeddings, incorporating temporal trends in lab values, or some extracts from the unstructured data, would likely improve performance. Finally, use of this APHRODITE-based pipeline relies on sites mapping their EHR data to the OMOP CDM.

## CONCLUSION

We demonstrate a high-throughput pipeline for generating high-precision phenotype classifiers by using APHRODITE with a high-precision labeling heuristic. These classifiers are easier to create than rule-based definitions and are portable across sites. We demonstrate good portability between Stanford and Columbia in both directions, but limited portability with SNUBH, suggesting that the generalizability of phenotype classifiers may have geographic limitations. Sharing the classifier training recipe – i.e the labeling function for generating a large imperfectly labeled training set and re-training classifier – showed higher portability in all comparisons suggesting that sharing the classifier-building recipe, rather than the pre-trained classifiers, may be more useful for facilitating collaborative observational research.

## REFERENCES

1 Banda JM, Seneviratne M, Hernandez-Boussard T, et al. Advances in Electronic Phenotyping: From Rule-Based Definitions to Machine Learning Models. Annu Rev Biomed Data Sci 2018;1:53–68. doi:10.1146/annurev-biodatasci-080917-013315

2 Richesson RL, Hammond WE, Nahm M, et al. Electronic health records based phenotyping in next-generation clinical trials: a perspective from the NIH Health Care Systems Collaboratory. J Am Med Inform Assoc 2013;20:e226–31. doi:10.1136/amiajnl-2013-001926

3 Hripcsak G, Albers DJ. Next-generation phenotyping of electronic health records. J Am Med Inform Assoc 2013;20:117–21. doi:10.1136/amiajnl-2012-001145

4 Pathak J, Kho AN, Denny JC. Electronic health records-driven phenotyping: challenges, recent advances, and perspectives. J Am Med Inform Assoc 2013;20:e206–11. doi:10.1136/amiajnl-2013-002428

5 Shivade C, Raghavan P, Fosler-Lussier E, et al. A review of approaches to identifying patient phenotype cohorts using electronic health records. J Am Med Inform Assoc 2014;21:221–30. doi:10.1136/amiajnl-2013-001935

6 Kirby JC, Speltz P, Rasmussen L V, et al. PheKB: a catalog and workflow for creating electronic phenotype algorithms for transportability. J Am Med Inform Assoc 2016;23:1046–52. doi:10.1093/jamia/ocv202

7 Newton KM, Peissig PL, Kho AN, et al. Validation of electronic medical record-based phenotyping algorithms: results and lessons learned from the eMERGE network. J Am Med Inform Assoc 2013;20:e147–54. doi:10.1136/amiajnl-2012-000896

8 Hripcsak G, Duke JD, Shah NH, et al. Observational Health Data Sciences and Informatics (OHDSI): Opportunities for Observational Researchers. Stud Heal Technol Inform 2015;216:574–8. https://www.ncbi.nlm.nih.gov/pubmed/26262116

9 Kohane IS, Churchill SE, Murphy SN. A translational engine at the national scale: informatics for integrating biology and the bedside. J Am Med Inform Assoc 2012;19:181–5. doi:10.1136/amiajnl-2011-000492

10 Tian TY, Zlateva I, Anderson DR. Using electronic health records data to identify patients with chronic pain in a primary care setting. J Am Med Inform Assoc 2013;20:e275–80. doi:10.1136/amiajnl-2013-001856

11 Carroll RJ, Eyler AE, Denny JC. Naïve Electronic Health Record phenotype identification for Rheumatoid arthritis. AMIA Annu Symp Proc 2011;2011: 189–96. https://www.ncbi.nlm.nih.gov/pubmed/22195070

12 Carroll RJ, Thompson WK, Eyler AE, et al. Portability of an algorithm to identify rheumatoid arthritis in electronic health records. J Am Med Inform Assoc 2012;19:e162–9. doi:10.1136/amiajnl-2011-000583

13 Chen Y, Carroll RJ, Hinz ERM, et al. Applying active learning to high-throughput phenotyping algorithms for electronic health records data. J Am Med Inform Assoc 2013;20:e253–9. doi:10.1136/amiajnl-2013-001945

14 Agarwal V, Podchiyska T, Banda JM, et al. Learning statistical models of phenotypes using noisy labeled training data. J Am Med Inform Assoc 2016;23:1166–73. doi: 10.1093/jamia/ocw028

15 Halpern Y, Choi Y, Horng S, et al. Using Anchors to Estimate Clinical State without Labeled Data. AMIA Annu Symp Proc 2014;2014:606–15. https://www.ncbi.nlm.nih.gov/pubmed/25954366

16 Halpern Y, Horng S, Choi Y, et al. Electronic medical record phenotyping using the anchor and learn framework. J Am Med Inform Assoc 2016;23:731–40. doi: 10.1093/jamia/ocw011

17 Beaulieu-Jones BK, Greene CS, Consortium PRO-AALSCT. Semi-supervised learning of the electronic health record for phenotype stratification. J Biomed Inform 2016;64:168–78. doi: 10.1016/j.jbi.2016.10.007

18 Yu S, Chakrabortty A, Liao KP, et al. Surrogate-assisted feature extraction for high-throughput phenotyping. J Am Med Inform Assoc 2017;24:e143–9. doi:10.1093/jamia/ocw135

19 Murray SG, Avati A, Schmajuk G, et al. Automated and flexible identification of complex disease: building a model for systemic lupus erythematosus using noisy labeling. J Am Med Informatics Assoc 2019;26:61–5. doi:10.1093/jamia/ocy154

20 Simon HU. General Bounds on the Number of Examples Needed for Learning Probabilistic Concepts. J Comput Syst Sci 1996;52:239–54. doi:10.1006/jcss.1996.0019

21 Banda JM, Halpern Y, Sontag D, et al. Electronic phenotyping with APHRODITE and the Observational Health Sciences and Informatics (OHDSI) data network. AMIA Jt Summits Transl Sci Proc 2017;2017:48–57. https://www.ncbi.nlm.nih.gov/pubmed/28815104

22 Pacheco JA, Rasmussen L V, Kiefer RC, et al. A case study evaluating the portability of an executable computable phenotype algorithm across multiple institutions and electronic health record environments. J Am Med Inform Assoc 2018;25:1540–6. doi:10.1093/jamia/ocy101

