## Supplementary material for "Development and Validation of Phenotype Classifiers across Multiple Sites in the Observational Health Sciences and Informatics (OHDSI) Network"

| Phenotype | Characteristic | Stanford | Columbia | SNUBH | p value* |
|---|---|---|---|---|---|
| Appendicitis | Proportion of males | 0.52 | 0.52 | 0.51 | 0.98 |
| | Mean age | 34.8 | 34.14 | 44.49 | |
| T2DM | Proportion of males | 0.51 | 0.44 | 0.56 | < 0.05 |
| | Mean age | 63.61 | 67.77 | 66.47 | |
| Cataracts | Proportion of males | 0.45 | 0.37 | 0.43 | < 0.05 |
| | Mean age | 68.84 | 75.54 | 68.22 | |
| HF | Proportion of males | 0.56 | 0.50 | 0.48 | < 0.05 |
| | Mean age | 65.24 | 70.69 | 72.9 | |
| AAA | Proportion of males | 0.77 | 0.68 | 0.79 | 0.56 |
| | Mean age | 75.5 | 78.24 | 77.41 | |
| Epileptic seizure | Proportion of males | 0.55 | 0.47 | 0.53 | 0.30 |
| | Mean age | 38.44 | 45.57 | 32.27 | |
| PAD | Proportion of males | 0.6 | 0.48* | 0.69 | < 0.05 |
| | Mean age | 71.81 | 73.29 | 66.22 | |
| Adult onset obesity | Proportion of males | 0.35 | 0.30* | 0.61* | < 0.05 |
| | Mean age | 53.4 | 45.18 | 35.9 | |
| Glaucoma | Proportion of males | 0.47 | 0.40 | 0.58 | < 0.05 |
| | Mean age | 72.1 | 80.10 | 62.97 | |
| VTE | Proportion of males | 0.55 | 0.39 | 0.51 | < 0.05 |
| | Mean age | 62.26 | 65.02 | 68.46 | |
| **All patients** | **Proportion of males** | **0.45** | **0.45** | **0.48** | **< 0.05** |
| | **Mean age** | **39.30** | **39.90** | **40.41** | |

**Supplemental Table 1.** Comparison of key demographics in patients identified by phenotype classifiers at three sites within OHDSI network. p value for comparing proportion of males was calculated using chi squared test.
